# Loss of O-specific antigen shapes *Pseudomonas aeruginosa* population microbiogeography in murine preclinical pulmonary infection model

**DOI:** 10.1101/2024.10.21.619428

**Authors:** H. Fraser, D. A. Moustafa, J. B. Goldberg, S. Azimi

## Abstract

Chronic *Pseudomonas aeruginosa* infections are the hallmark of late-stage lung disease in individuals with cystic fibrosis. During chronic infection *P. aeruginosa* becomes the dominant bacteria in the airway. Within-host adaptation of *P. aeruginosa* leads to vast phenotypic and genetic population heterogeneity. *In vitro* studies show mutations in lipopolysaccharide (LPS) O-specific antigen changes the aggregate formation in *P. aeruginosa*, however role of these changes in aggregate assembly *in vivo* is not understood. Using a synthetic CF sputum media and a preclinical murine infection model we assessed how the PAO1 wildtype and O-specific antigen mutants interact with each other, and if *P. aeruginosa* population heterogeneity affects the colonization of the murine lungs. Our findings suggest that the presence of variants lacking O-specific antigen does not impact the population fitness and size in both *in vitro* and *in vivo*, however it can influence the aggregate volume *in vivo*.

---

Chronic infections are usually polymicrobial, where bacteria live in biofilms or as suspended aggregates of 10–10^4^ cells with defined spatial patterning and distances relative to host cells and other microorganism^1-3^. In chronic infections, within-host adaptation, genetic drift, and antimicrobial selection are key drivers of intrastrain genetic heterogeneity of bacterial populations. Although the intrastrain genetic heterogeneity is known, its impact on pathogenesis, and microbiogeography of infections remains unclear.

In chronic pulmonary infection with *Pseudomonas aeruginosa* in people with cystic fibrosis (pwCF) is the hallmark of loss of lung function. Divergent evolution of *P. aeruginosa*, and accumulation of different genetic variants in CF airways and its impact on population functional phenotypes is well documented^3-6^, but how this intrastrain heterogeneity influences polymicrobial community and airway focal injury remains unknown. We recently showed that mutations in genes encoding Lipopolysaccharide (LPS) O-specific antigen (OSA) biosynthesis and assembly such as *ssg* and *wbpL* increase *P. aeruginosa* cell surface hydrophobicity, and alter *P. aeruginosa* aggregate assembly in *in vitro* preclinical CF sputum model (SCFM2)^7-9^. Considering *P. aeruginosa* variants with mutations in OSA commonly isolated from chronic CF airway infections^10-13^, along with LPS smooth phenotypes, it is important to determine the role of OSA genetic and phenotypic heterogeneity on spatial organization, and *P. aeruginosa* population fitness.

To address the influence of genetic heterogeneity on spatial organization of *P. aeruginosa* population in the airways, we examined whether presence of variants with mutations in OSA biosynthesis and assembly influence the aggregate assembly of *P. aeruginosa* populations *in vitro*. We constructed mixed populations of PAO1WT with either PAO1Δ*ssg* or, PAO1Δ*wbpL* at 1:1 initial ratio (5×10^6^: 5× 10^6^ CFU) in SCFM2. We grew each variant and the parental PAO1 strain in isolation and mixed populations for 24 hours and evaluated the changes in total population size, and changes in the ratio of PAO1Δ*ssg* and PAO1Δ*wbpL* compared to the wild type PAO1 strain. We assessed the abundance of each variant by qPCR and by enumerating colony forming unites (CFUs). Although we detected a significant change in aggregate assembly of WT cells in presence of ΔOSA variants (PAO1Δ*ssg* and PAO1Δ*wbpL*), we did not detect any significant changes in proportion of each variant in mixed populations *in vitro* (Fig. 1 and b). Interestingly, we observed that presence of ΔOSA variants significantly disrupts the stacked aggregate assembly of parental strain (Fig. 1c).

**Fig. 1.**
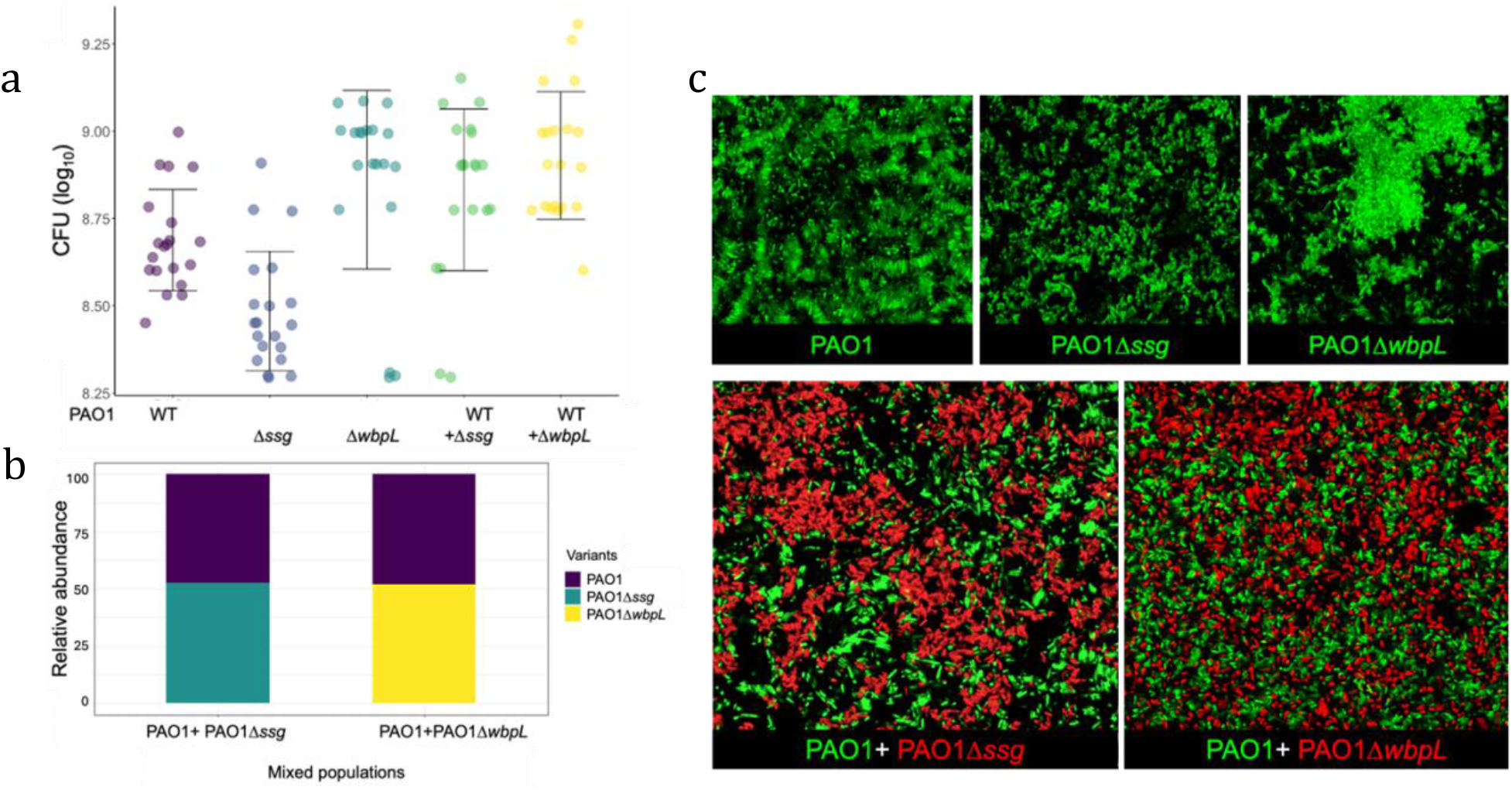
Loss of OSA due to mutations in *ssg* and *wbpL* genes, alters the biogeography and aggregate assembly of *P. aeruginosa* populations. (a) Presence of both PAO1Δ*ssg* and PAO1Δ*wbpL* lead to increase of total population size, but (b) not the population structure, as the relative abundance of PAO1, PAO1Δ*ssg* and, PAO1Δ*wbpL* in mixed populations remains the same. (c) Changes of aggregate assembly from Stacked to Clumped due to loss of OSA and increases cell surface hydrophobicity, and presence of variants without OSA alters the biogeography of mixed populations (lower panel), (data presented from three independent experiments).

Although presence of ΔOSA variants alters the population biogeography *in vitro*, it is not clear how the changes in population structure affect the spatial organization in airways. To examine how presence of ΔOSA variants influence airway colonization, we modified an established preclinical acute airway infection model^14, 15^ of six-week-old female BALB/c mice. We prepared cultures of PAO1, PAO1Δ*ssg*, PAO1Δ*wbpL*, and synthetic mixed populations of PAO1: PAO1Δ*ssg* and PAO1: PAO1Δ*wbpL* in SCFM2 and incubated the cultures 4-6 hours at 37 °C. We then infected the mice with 25 μL of the standard inoculum 1x 10^7^ CFU of each bacterial population. We then sacrificed the mice 24 hours post infection for further analysis of colonization rate and bacterial spatial organization in the airways (All animal procedures were conducted according to the guidelines of the Emory University Institutional Animal Care and Use Committee, under approved protocol number PROTO201700441). Briefly, we determined the abundance of each *P. aeruginosa* variants by homogenizing the lungs, and enumerating CFUs^15^, and qPCR (Fig. 2 a and b). In brief, we perfused the lungs with 15 ml of 20Uml of heparin in PBS. We then fixed the lungs in 500 ml of 4% PFA/PBS (v/v) for future immunohistochemistry, hybridization chain reaction (HCR)-probing^16, 17^, and tissue-clearing following iDISCO^18^(Fig. 2c).

**Fig. 2.**
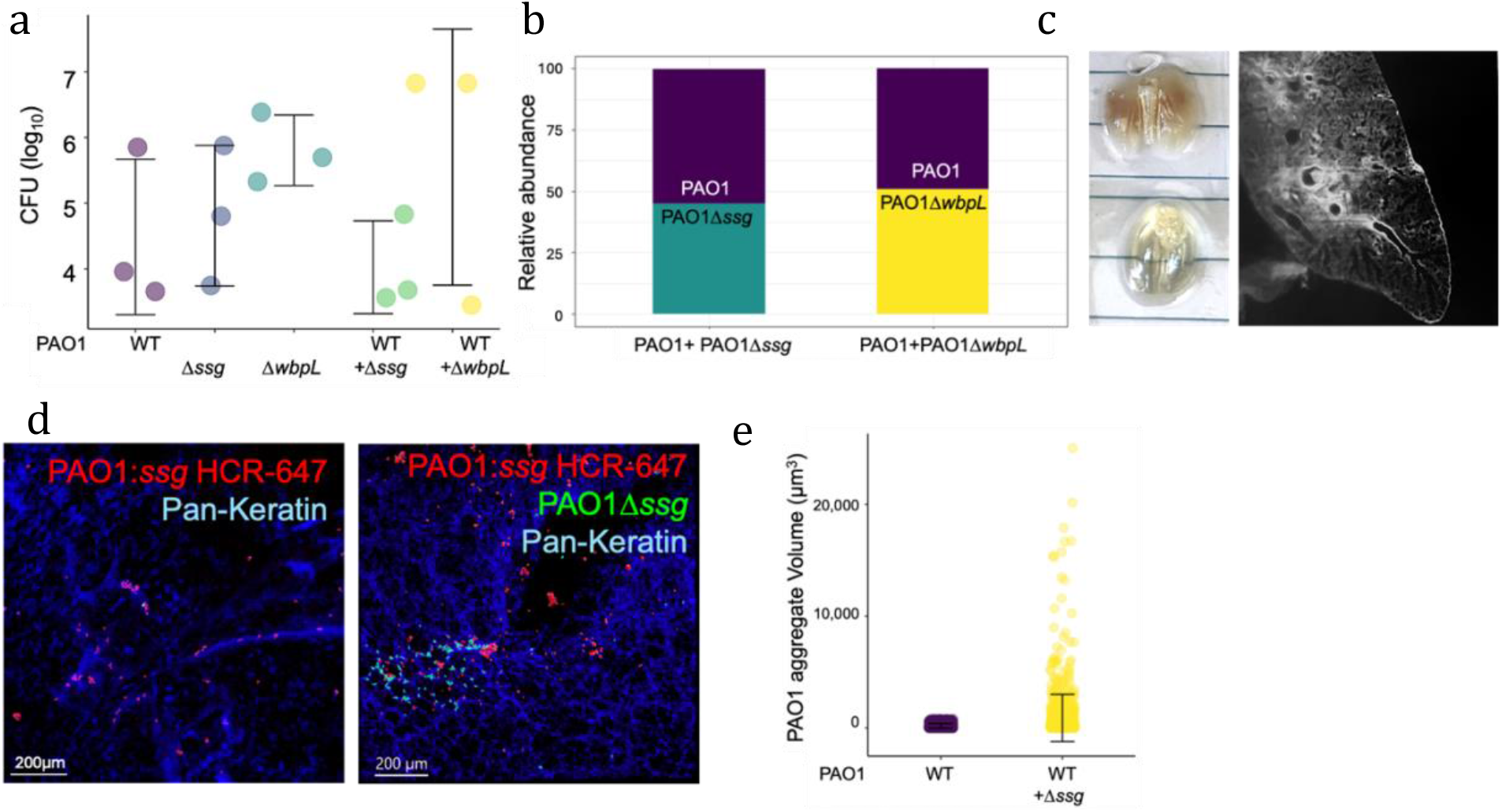
Presence of PAO1Δ*ssg* increase the PAO1 aggregate size in murine airway. (a) Presence of ΔOSA variants in a population does not significantly impact colonization and population size in murine airways. (b) Loss of OSA does not influence *P. aeruginosa* fitness in murine airway. (c) Use of iDISCO clearing method allows for whole lung imaging and localization of bacteria in the airways. (d) Presence of PAO1Δ*ssg* does not affect the abundance of WT cells but (e) leads to formation of larger aggregates (df= 1011.2, *p*< 0.001).

We measured the colonization level of each variant and mixed populations in the airways by homogenizing the lungs in 1ml of PBS, followed by enumerating the CFUs. We further extracted the genomic DNA from lung homogenates and measured the abundance of each variant in mixed populations using qPCR. We found that similar to what we observed in *in vitro*, loss of OSA due to deletion of *ssg* and *wbpL* does not influence *P. aeruginosa* fitness in murine airways either in single or in mixed populations (Fig. 2 b and c). To determine whether presence of ΔOSA variants in *P. aeruginosa* populations alter the spatial organization and microbiogeography of the airways, we used confocal and light sheet microscopy. To visualize the bacteria in the lungs, we infected the mice with PAO1Δ*ssg*: *gfp* and used HCR-probes targeting *ssg* mRNA to detect and visualized PAO1 WT cells. We used Imaris.10 software to measure the aggregate volumes and observed a significant change in PAO1WT aggregate volumes in presence of PAO1Δ*ssg* cells from average 178 μm^3^ to 898 μm^3^ (Fig. 2 d and e).

In summary, our study highlights the significant role that genetic heterogeneity within *P. aeruginosa* populations plays in shaping bacterial spatial organization, particularly in chronic infections such as those found in CF. Despite no measurable impact on overall population fitness, the presence of variants with OSA mutations notably alters aggregate assembly both *in vitro* and *in vivo*. These findings suggest that population genetic heterogeneity have the potential to influence the structural organization of bacterial communities, which can subsequently impact the infection progression, host immune responses at micron scale, and treatment outcomes. Future studies assessing the impacts of genetic heterogeneity in clinically sourced populations, on host pathogen interactions and pathogenesis, will be crucial for understanding the broader implications of *P. aeruginosa* genetic diversity in chronic infections and focal airway injury.

## Acknowledgement

We would like to acknowledge members of the Azimi, and Goldberg labs for their assistance in setting up the animal experiments. This study was supported by Georgia State University startup to SA, and CFF-WHITELE20A0 to JBG.

