## Supplemental methods for "Loss of O-specific antigen shapes *Pseudomonas aeruginosa* population microbiogeography in murine preclinical pulmonary infection model"

#### Bacterial strains and growth conditions.

We used PAO1strain of *P. aeruginosa*, and isogenic mutants PAO1Δ*ssg,* and PAO1Δ*wbpL* with ΔOSA phenotype. For all *in vitro* experiments in SCFM2[1,2], we grew the bacterial strains in 3 ml of lysogeny broth (LB) to mid-log phase at 37˚C/200 *rpm*. We then measured the $OD_{600}$ and adjusted the bacterial density to $OD_{600}$ = 0.01 in 3 ml of SCFM2 and incubated for 4-6 hours for either animal experiments or $OD_{600}$ = 0.01 in 400 μl of SCFM2 for imaging; static at 37˚C. For each experiment, we prepared fresh SCFM2 as described[1]. Briefly, we prepared a 50 ml buffered base stock solution containing basal salts and amino acids, that was adjusted to a pH of 6.8 using hydrochloric acid and then filter sterilized. The day before the SCFM2 was needed we prepared 50 ml by adding 0.6 mg/ml salmon sperm DNA (SIGMA) and 5mg/ml porcine maxillary mucin (SIGMA). On the day the SCFM2 was used we added 500 μl each of dextrose, L-lactic acid, calcium chloride, and magnesium chloride, fresh iron-sulphate heptahydrate, N-acetylglucosamine and 1,2- dioleoyl-sn-glycero-3-phosphocholine.

#### Animal Infection Model

We grew PAO1 in LB broth for 16-18 h at 37˚C and then suspended it in PBS to an adjusted $OD_{600}$ of 0.05, corresponding to ~$10^7$ CFU/ml in SCFM2, and incubated for an additional 6 hours at 37˚C for aggregate assembly. We adjusted to obtain the desired challenge dose in a volume of 50 μL. Six-week-old female Balb/c mice (Jackson Laboratories, Bar Harbor, ME). We infected the mice by non-invasive intratracheal instillation of dilutions of each mixed population and single *P. aeruginosa* (WT, PAO1Δ*ssg,* and PAO1Δ*wbpL*). We euthanized the mice at 24 h post-infection and either collected whole lungs aseptically, weighed, and homogenized in 1 mL of PBS for serially dilution and plating on PIA and CFU determinations[3], or perfused the lungs and then fixed the lungs in 4% PFA/PBS (w/v) for further tissue clearing. All animal procedures were conducted according to the guidelines of the Emory University Institutional Animal Care and Use Committee, under approved protocol number DAR-2003421–042219BN.

#### Quantification of *in vitro* and *in vivo* bacterial aggregate biomass

For *in vitro* assays*,* we grew each strain to mid-log phase in LB at 37˚C/200 *rpm*. We then measured the $OD_{600}$ and adjusted the bacterial density to $OD_{600}$ = 0.01 in 3 ml of SCFM2, and incubated overnight, static at 37˚C. For mixed populations, a 1:1 ratio of PAO1 WT to PAO1 LPS mutant strains to final $OD_{600}$ = 0.01 ($OD_{600}$ = 0.005 for each variant) starting cells in each culture.
For the *in vivo* assays we homogenized the harvested lung from each of the mice using a bead beater in 1ml of PBS. We used 200 μl of each culture to measure the final CFU of each population. We added 40 μl of 5% Tween-20 (final concentration of 1%) to break down bacterial aggregates, followed up by serially diluting the cultures with PBS (1:10) and inoculating 5 μl of each dilution to LB agar plates.

#### Bacterial abundance analysis using qPCR

We extracted the DNA from 1 ml of overnight bacterial cultures in SCFM2 using Wizard Genomic DNA purification kit (Promega) following the manufacturer's protocol. We used the nanodrop to measure the DNA concentration extracted from each sample. For murine samples we used DNeasy Blood and Tissue kit (Qiagen) to extract total DNA from lung homogenates.

For qPCR, we used SYBR green PCR mix (Applied Biosystems). For each experiment, both *in vitro* and *in vivo*, we diluted all DNA samples to 50 ng/µl so and used 250 ng/µl for each reaction. We designed and used three sets of primers for *gyrA*, *ssg,* and *wbpL* to detect abundance of *P. aeruginosa* (Table. 1).

| | |
|---|---|
| *ssg* F | TTTTCCCGCTCGCCCTGGTCGT |
| *ssg* R | TGCTCGACGAGTTGGGCGGAGT |
| *wbpL* F | GGCGTGGCACTGTTAGGGTTCC |
| *wbpL* R | TTCCAGATCAGGAAGCCGGCGA |
| *gyrA* F | TGTGCTTTATGCCATGAGACGA |
| *gyrA* R | CGTTGAAGCCAAGCCACCT |

### Tissue Clearing methods

#### iDISCO tissue clearing

We followed the optimized iDISCO protocol[4]. Briefly, we dehydrated the tissue by incubating it in 20%, 40%, 60%, 80%, and 100% methanol in water washes for 1 hour each. Then we incubated the tissue in DCM/Methanol 2:1 for 3 hours at room temperature, shaking. We then washed the tissues twice with DCM for 15 minutes. Finally, the samples were incubated in DiBenzyl Ether (SIGMA) until the samples became transparent.

#### Hybridization Chain Reaction Fluorescent *in situ* hybridization (HCR-FISH)

We blocked the tissues in 4% (v/v) bovine serum albumin/Tris-Buffered Saline (BSA/TBS) solution for three days at room temperature shaking. We used specific HCR v3.0[5] probes (Molecular Instruments) to detect different variants in the infected lung tissues. We tested the specificity of HCR probes in *in vitro* cultures of *P. aeruginosa* to ensure the probes would bind to the correct mRNA sequence following the manufacturer protocol[5].

#### Immunofluorescent staining and

We added the primary antibody, anti-Mouse pan Keratin IgG (Abcam), to 4% BSA at 1:1000 dilution and incubated for 1 hour at room temperature. We then washed the tissues once with 0.01% (v/v) of PBS-Tween 20 for 15 minutes, followed by three washes with PBS for 15 minutes. Next, we incubated the tissues with Goat Anti-Rabbit IgG H&L at 1:80 dilution overnight at 4°C followed by PBS-Tween 20 washes. We stored the tissues at 4% BSA/TBS solutions at 4°C.

#### Murine tissue image acquisition and analysis

To image the tissue samples, we used an LSM880 confocal laser scanning microscope (Zeiss)., we used Imaris 10.0.1 image analysis software. We measured the volume of the aggregates in 6 different images for each condition (over 900 aggregates were measured in total per each condition).
